# Neural Architectures of Slow and Fast Dynamics in the Human Brain

**DOI:** 10.64898/2025.12.31.696728

**Authors:** Yuhan Lu, Hangze Mao, Zhuoran Li, Qinsiyuan Lyu, Yiduo Lu, Chen Yao, Junxi Chen, Louis Tao, Zhuo-Cheng Xiao, Xing Tian

## Abstract

The human brain operates across a vast temporal range, from fast perception and action to slow physiological regulation. The capacity has attributed to a unitary cortical gradient of intrinsic timescales, yet such a unidimensional model cannot explain how local circuits simulateously support both rapid external behavior and slow internal body-coupled dynamics. Using SPLIT (<u>s</u>pectral <u>p</u>iecewise-linear inference of timescales) on a large-scale intracranial stereo-electroencephalography (8,619 contacts, 185 individuals), we identified dissociable fast (∼10-100 Hz) and slow (∼1-10 Hz) temporal components. Only the fast-component timescales followed the canonical sensorimotor-to-association cortical hierarchy. Slow-component timescales showed no hierarchical gradient, but were instead covaried with heart rate and enriched at site with heart-related neural responses. This dual temporal architecture persisted across wakefulness, rest, sleep, and anesthesia, challenging the unitary view of brain timescales and revealing an intrinsic bipartite organization that is associated with cortical hierarchy or neurophysiology.

**Significance statement:** The brain’s temporal organization is commonly described by a hierarchy in which sensory regions operate rapidly and association regions integrate information over longer periods. We show that this account captures only part of human neural dynamics. Intracranial recordings from 185 individuals reveal dissociable fast and slow components, often coexisting locally. Only the fast component follows the cortical hierarchy, whereas the slow component is associated with neural responses to heartbeats. This dual-component organization persists across wakefulness, rest, sleep, and anesthesia. The findings extend the brain’s temporal architecture beyond a single cortical gradient to include a distinct component linked to bodily physiology.

## Introduction

To survive and flourish, the human brain must rapidly interact with the environment while maintaining stable internal physiology. Therefore, the brain must support computations across orders of magnitude in time, ranging from rapid sensory processing and action to slow cognitive integration and allostatic regulation (1–5). A prevailing view holds that this multiscale capacity is unidimensionally embedded in a cortical hierarchy, characterized by a monotonic increase of the intrinsic neural timescale from unimodal to transmodal regions (6–12) (i.e., the gradient of timescales). However, this hierarchy-spanning network architecture is also implicated in much slower interoceptive monitoring and visceromotor control of peripheral physiology (13, 14). This raises a key organizational question: does each region operate around a single dominant intrinsic timescale, such that the cortex is described by one unidimensional gradient, or do local circuits contain separable temporal components that map onto multiple, parallel gradients serving distinct functional demands (**Fig. 1a**)? Investigating the organizing principle would provide a holistic picture of the brain’s temporal architecture, the scaffold on which neural computations unfold (8, 15, 16) and whose alteration drives transitions from healthy cognition to neuropsychiatric disorders (17, 18).

**Figure 1.**
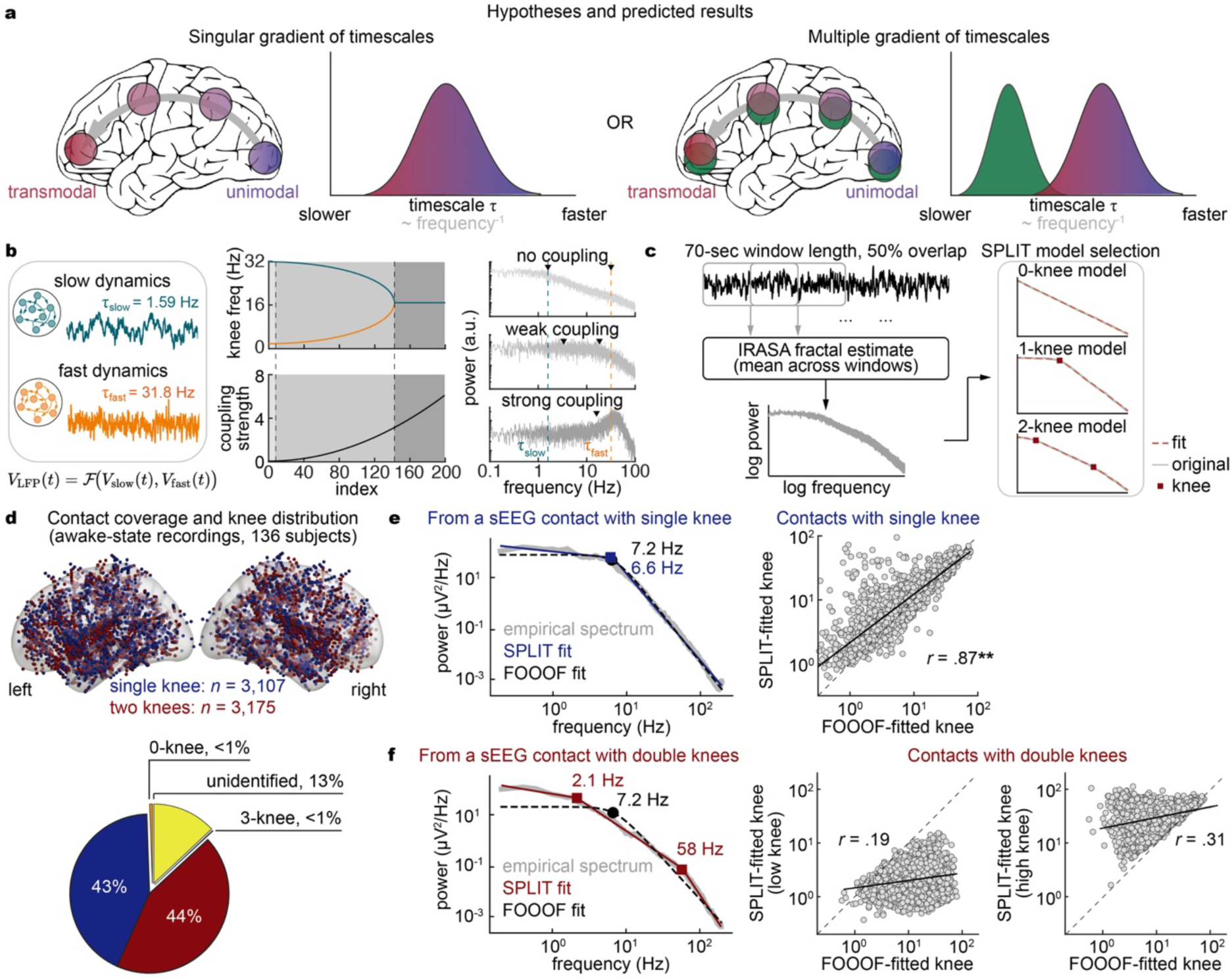
Competing hypotheses for the cortical gradient of neural timescales, and assumption-based timescale estimation in human sEEG. (**a**) Hypotheses about the gradient of timescales. Left: a dominant timescale in local regions comprises a single gradient of timescales across the brain hierarchy, increasing from unimodal to transmodal cortex. Right: multiple concurrent local timescales across the brain hierarchy. (**b**) Analytical model motivating timescale estimation without assuming its number. Left, the LFP is modeled as a mixture of two Ornstein-Uhlenbeck populations with distinct time constants (i.e., expressed as 1.59-Hz and 31.8-Hz knees). Middle, knee frequencies of the resulting spectrum (top) as a function of inter-population coupling strength (bottom) across 200 parameter settings; beyond a critical coupling (dashed line, dark shading) the two knees merge into one. Right, representative spectra under no, weak, and strong coupling; arrowheads mark fitted knees and dashed lines the ground-truth f_slow_ (blue) and f_fast_ (orange). Uncoupled populations preserve two knees; strong coupling collapses the dynamics onto a single eigenmode and one knee. (**c**) SPLIT estimation pipeline. Power spectra were computed in 70-s windows with 50% overlap; the aperiodic (fractal) component was isolated by IRASA and averaged across windows. Models with 0, 1, or 2 knees (red squares, fits in dashed red) were fitted to the log-log spectrum, without assuming the number of timescales a priori. (**d**) Top, sEEG contact coverage during awake rest (136 patients), colored by the best-fitting model (blue, single knee, *n* = 3,107; red, two knees, *n* = 3,175). Bottom, proportion of contacts assigned to each model. (**e**) Contacts with single-knee spectra. Left, spectrum from a representative contact (gray) fitted by SPLIT (blue, knee at 7.2 Hz) and by FOOOF, which assumes one knee (black dashed, 6.6 Hz). Right, across all single-knee contacts (*n* = 3,107), knee frequencies from the two methods agree closely. (**f**) Contacts with double-knee spectra. Left, spectrum from a representative contact; SPLIT resolves two knees (red, 2.1 and 58 Hz), whereas the single-knee assumption forces FOOOF to a single intermediate value (black, 7.2 Hz). Right, across all double-knee contacts (*n* = 3,175), the FOOOF estimate correlates only weakly with the SPLIT low (*r* = 0.19) and high (*r* = 0.31) knee frequencies, indicating that imposing one knee neither recovers either timescale nor flags the mismatch. \*\**p* < 0.05.

Addressing this organizational question requires revisiting a tacit assumption that has guided most work on the brain-wide temporal hierarchy: a region’s activity can be approximated by a single dominant timescale. Accordingly, widely-used methods, such as autocorrelation decay (9, 10, 19–24) or spectral knee frequency (25–29), reduce an aggregated time series (e.g., local field potentials [LFPs] or BOLD signals) to a single timescale per region, and the “gradient of timescale” is inferred by comparing these regional scalars across cortices (10, 25, 30). However, multiple timescales can co-exist within the same local circuits (28, 29, 31, 32), making single-timescale reduction oversimplified and potentially biased because it conflates distinct temporal components, and therefore challenging the unidimensional view of the brain’s temporal architecture.

Resolving the debate about the organizing principle of brain-wide temporal infrastructure requires recordings with millisecond temporal precision and broad anatomical coverage (3, 9, 32), which is an uncommon combination in human neuroscience. Large-scale intracranial stereo-electroencephalography (sEEG) bridges this gap by sampling mesoscopic population dynamics at high rates across cortex and deep structures, providing a natural substrate for linking microcircuit timescale diversity to brain-wide organization. We therefore approach the question as component-resolved inference: by identifying timescales within LFPs without a single-timescale prior, we examine the brain-wide distribution of timescales and examine whether the human brain’s temporal infrastructure follows a unidimensional gradient or dissociable architectures.

Applying the SPLIT (<u>s</u>pectral <u>p</u>iecewise-linear inference of timescales) algorithm for robust estimation of timescales to a large-scale human sEEG recordings (total of 8,619 contacts from 185 subjects), we revealed a brain-wide organization of dissociable fast (∼10-100 Hz) and slow (∼1-10 Hz) components of timescales. Only the fast component recapitulated the canonical cortical hierarchy, whereas the slow component was ubiquitously related to heart-related neural responses, with no detectable cardiac field leakage. The two-track brain-wide temporal infrastructure was stable across different conscious states, demonstrating the inherent nature of the neural architecture. The human brain operates on parallel temporal infrastructures for collectively regulating the bodily apparatus and interacting with the environment.

## Results

### The number of timescales cannot be fixed in advance

Empirical spectra may take a range of forms, from a 1/f-like decay to one or several bends, but how these shapes mapped onto the timescales of the underlying circuit was not obvious. To understand what determined which of these appears, we simulated a minimal two-dimensional Ornstein-Uhlenbeck (OU) system with a fast and a slow time constant and examined the knees in the resulting spectra (**Fig. 1b**). Hereafter we reported spectral knees in Hz to express timescales, the two being related by τ = 1/(2π*f*_knee_). When the two processes evolved independently, both knees appeared at the frequencies set by their own time constants (uncoupling condition). When we allowed the two processes to interact, the knees shifted away from their individual time constants and instead followed the combined system (weak coupling condition), and above a certain interaction strength they merged into one, leaving a single knee with a narrowband peak beside it (strong coupling condition). Therefore, whether a spectrum expressed one or two knees depended on the number of underlying processes and on how strongly they interacted, neither of which was known before the spectrum was fitted. Estimating timescales from a spectrum therefore required leaving the number of knees free rather than fixing it in advance.

### SPLIT inferred the number and values of timescales from the power spectrum

We therefore developed a spectral analysis procedure without imposing a prior on the number of timescales (i.e., SPLIT; see Methods for full details) that inferred the number and frequency values of the spectral knees directly from the PSD. The method exploited the fact that distinct timescales manifested as successive linear-like regimes in the log-log PSD (8, 27), with knees marking transitions between approximately linear segments (25). For each contact, the aperiodic spectrum was isolated by IRASA in overlapping windows and averaged across windows, and the averaged spectrum was submitted to the SPLIT fit (**Fig. 1c**). Based on OU simulations, we showed that low-frequency knees were systematically overestimated at short window lengths and, to recover knees down to ∼0.2 Hz within 20% error, we used 70-s windows throughout (**Fig. S1**). We also selected IRASA to separate aperiodic spectrum because it well preserved the fractal component and the knees within it (**Fig. S2**). Spectra were fitted over 0.2 to 200 Hz, a range encompassing the knee frequencies previously reported in LFP recordings (e.g., ∼0.1-10 Hz (28), ∼75 Hz (29), and also see (32–34)). Knees below this range (∼0.01 Hz, equivalent to second-level timescale) require both long recordings and a sufficiently low hardware high-pass, and may be the product of artifacts; they were outside the scope of the present analysis (**Fig. S3**).

We benchmarked SPLIT using simulations with known knee numbers and frequencies, including idealized piecewise-linear spectra with additive noise, signals generated from one or two OU processes, and OU signals with superimposed oscillations. In the idealized spectra, the recovered knee number closely matched ground truth, and knee-frequency errors remained small across the tested noise levels (**Fig. S4a-b**). In OU signals, estimation accuracy depended on window duration and knee configuration: lower-frequency knees required longer windows, and the low-frequency knee in two-process mixtures was estimated less precisely than the high-frequency knee (**Fig. S4c-d**). We also showed that the position of oscillation relative to spectral knees affected the fitting error in the one-knee spectra and two-knee spectra (**Fig. S4e**). Together, these benchmarks supported the robustness of SPLIT in distinguishing one- and two-knee spectral classes.

Applying SPLIT to sEEG recordings acquired during wakefulness (e.g., subjects engaged in unstructured activity such as watching television or smart phone), most contacts showed one or two knees in the PSD between 0.2 and 200 Hz, with very few contacts showing zero or more than two (**Fig. 1d**). Approximately 13% of spectra were unidentified because their fitted segments did not show progressive steepening across successive knees, often accompanied by high-frequency flattening consistent with the noise floor. The one- and two-knee spectra were both prevalent and occurred at similar rates across the entire brain. This prevalence indicated that the assumption of a single timescale and the methods built on the assumption may potentially bias the fitting results and generated systematic errors. Next, we demonstrated this point by directly comparing a prevailing method (i.e., FOOOF (27)) that imposed a single-timescale assumption with the current method (i.e., SPLIT) that made no assumptions about the number of timescales.

### Systematic errors in fitting timescales by assuming a single timescale

For spectra identified as having one knee by SPLIT, the method that assumed a single-timescale and fitted by a single-Lorentzian function (FOOOF) produced highly correlated knee frequencies (*n* = 3,107; *r* = 0.87, Pearson’s *r* correlation; **Fig. 1e**). For two-knee spectra, by contrast, FOOOF-fitted knees corresponded only weakly to either the low- or the high-frequency knee identified by SPLIT (*n* = 3,175; *r* = 0.19 and 0.31, respectively; **Fig. 1f**), and fell systematically between them, above the low-frequency knee and below the high-frequency knee. Two accounts could explain this intermediate position (**Fig. S5a**). Under one, the spectrum contained a single knee that SPLIT overfits by inserting an additional segment, yielding two knees that straddled the true one recovered by FOOOF; under the other, the spectrum genuinely contained two knees, and FOOOF placed its single estimate between them because that position minimized the overall fitting error.

To distinguish them, we fitted the two-knee model to spectra that both methods had assigned a single knee (as shown in **Fig. 1e**) and examined the segment between the two fitted knees. In the single-knee spectra that the second linear segment of all three was forced across a gradual bend and should therefore fit poorly; if the two-knee spectra were falsely overfitted, then the second segment should be comparable with that in the single-knee spectra, but if it was itself a power law and should fit well (**Fig. S5b**). Comparing the two sets, the second linear segment was fitted poorer in the one-knee spectra (*p* = 0.0002, two-sided bootstrap, FDR corrected; **Fig. S5c**), and the two fitted knee frequencies were higher and lower, respectively, than those obtained from the two-knee spectra (*p* = 0.0002 for both comparisons; **Fig. S5c**). Therefore, the two-knee spectra resolved by SPLIT reflected genuine spectral structure rather than an overfitted single bend, and the intermediate value returned by FOOOF was a consequence of the single-timescale assumption.

### Knee frequencies clustered into fast- and slow-component timescales across the brain

To test whether the cortical temporal hierarchy was organized along a single axis or parallel axes (**Fig. 1a**), we examined the brain-wide distribution of all knee frequencies. Across both one-knee (*n* = 3,107; **Fig. 2a**) or two-knees contacts (low- and high-knee; *n* = 3,175; **Fig. 2b**), whose spectral knee frequencies did not form a single continuum but separated into a slow component at ∼1-10 Hz and a fast component at ∼10-100 Hz (**Fig. 2a-b**, right). A two-component Gaussian mixture described the single-knee distribution better than one- or three-component models (ΔBIC > 18.2 against both). The two components from single-knee spectra were separated at the intersection of the two Gaussians, and the similar two ranges naturally appeared in two-knee spectra, whose low and high knees were fitted within each spectrum without any clustering. Both components were present throughout the brain regions (**Fig. 2c**), supporting the multiple-gradient hypothesis.

**Figure 2.**
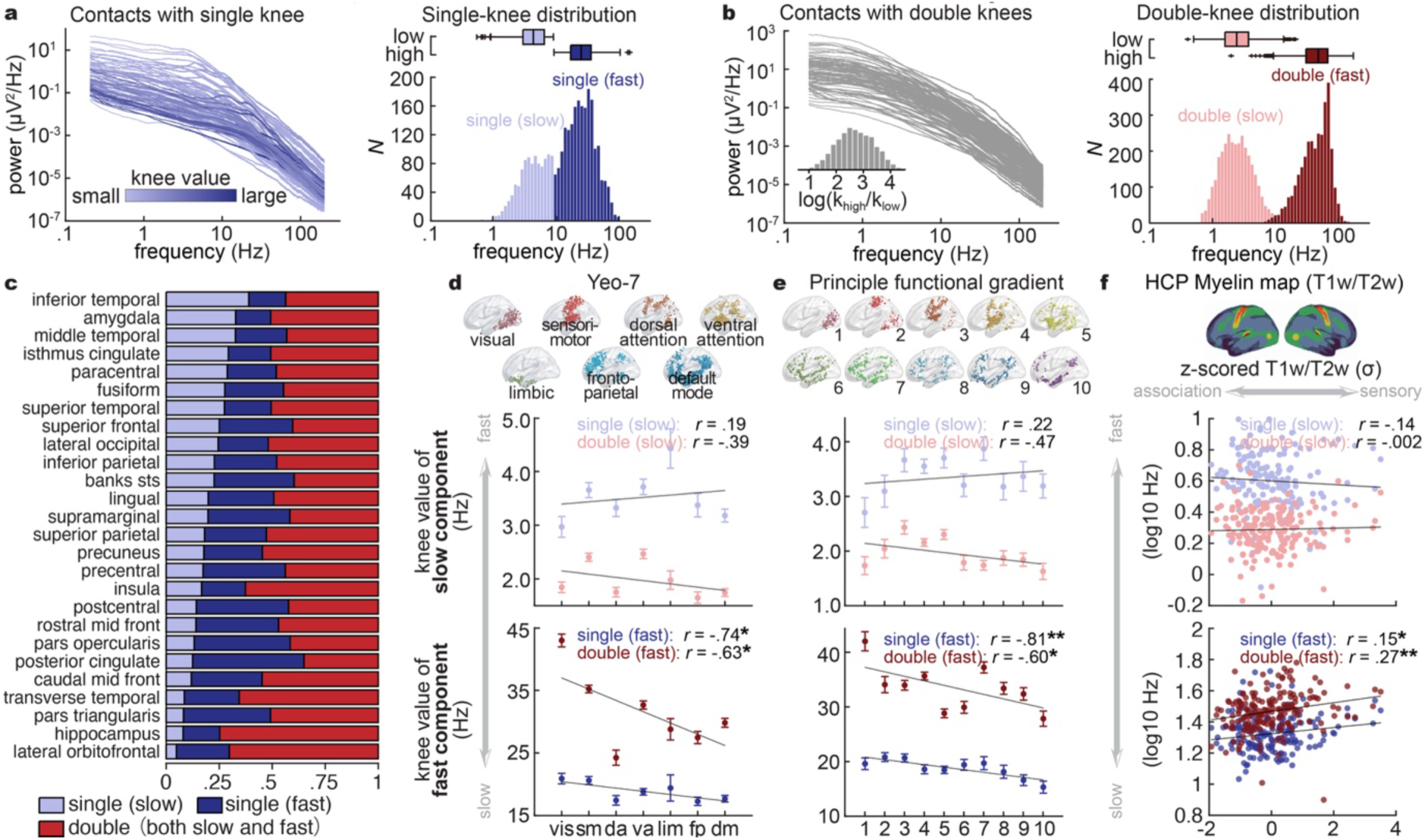
Two timescale components, of which only the fast one follows the cortical hierarchy. (**a**) Left: PSDs from single-knee contacts (*n* = 3,107). Each line represents PSD from a contact and is color-coded by the frequency value of the knee. Right: Distribution of knee frequencies for single-knee spectra. A bimodal profile was revealed, separating the distribution into slow and fast components. The inserted boxplots at the top summarize each component. (**b**) Left: PSDs from double-knee contacts (*n* = 3,175). Each line represents PSD from a contact. Insert, the ratio between the two knees follows a lognormal distribution centered at ∼2.5. Right: Distributions of low- and high-frequency knees for double-knee spectra. The bimodal distribution resembles similar frequency ranges observed in single-knee spectra in **a**. The slow and fast components naturally correspond to low- and high-knees, respectively. (**c**) Proportion of contact types across Desikan-Killiany regions. Bars give the proportion of contacts assigned to the slow (light blue) or fast (dark blue) component of the single-knee group, and to the double-knee group (red). (**d**) Frequency values of knee frequencies as a function of spatial segmentation in the Yeo-7 functional networks. Middle, the slow temporal component from both single- and double-knees spectra; Bottom, the fast temporal component from both single- and double-knees spectra. Only the fast temporal components correlate with the spatial gradient. Network abbreviations: vis, visual; sm, somatomotor; da, dorsal attention; va, ventral attention; lim, limbic; fp, frontoparietal; dm, default mode. (**e**) As in (**d**), across ten equally populated bins of the principal functional connectivity gradient, from unimodal/sensory (bin 1) to transmodal/association (bin 10). (**f**) Knee frequency against cortical myelination, an anatomical hierarchy proxy independent of functional connectivity. Top, group-average T1w/T2w ratio, z-scored; high values mark heavily myelinated sensory cortex, low values association cortex. Middle, slow component; bottom, fast component, with single-knee and double-knee contacts shown separately. Only the fast component tracked myelination, running faster toward sensory cortex. \**p* < 0.05, \*\**p* < 0.005.

### Only the fast-component timescale followed the cortical hierarchy

Crucially, these fast- and slow-component timescales were related to cortical hierarchy in different manners. Projected onto three established cortical annotations, i.e., the intrinsic functional gradients (35), the unimodal-transmodal principle gradient (36), and the myelin map (37) (**Fig. 2d-f**, top), the slow component showed no systematic variation across spatial regions (all *p* > 0.168, one-sided Pearson’s correlation, FDR corrected; **Fig. 2d-f**, middle). In contrast, the fast component exhibited a robust, monotonic gradient: the knee frequencies decreased from sensorimotor to association regions (all *p* < 0.035, one-sided Pearson’s correlation, FDR corrected; **Fig. 2d-f**, bottom). This pattern remained significant after accounting for inter-contact and inter-subject variance using linear mixed-effect models (fast component in both single- and double-knee spectrum: all *p* < 0.0007). Direct component-by-hierarchy tests further showed significant interactions across all three hierarchy measures for double-knee spectra (all *p* < 0.008), whereas interactions were marginal for single-knee spectra (all *p* < 0.076). Thus, the classical functional hierarchy was selectively embedded within the fast temporal component.

We further asked whether fitting the number of knees freely, as SPLIT did, was necessary for the separation into two components and for their dissociation along the hierarchy. We therefore refitted every spectrum with FOOOF (**Fig. S6**). For contacts that SPLIT had fitted with one knee, FOOOF recovered the same two components (**Fig. S6a**), as expected from **Fig. 1e**. For two-knee contacts (**Fig. S6b**) and for all contacts pooled (**Fig. S6c**), two components remained separable by BIC, but lay much closer together and were no longer distinguishable by inspection. Only when the distributions were explicitly separated into two components, all three showed that the fast component tracking every hierarchy measure and the slow component none. In contrast, pooling knee values without separation weakened the hierarchical relationship below significance for both functional measures and halved it for myelination (**Fig. S6d** versus **S6c**). Therefore, FOOOF can reproduce the previously reported cortical gradient when the fitting range is restricted to isolate the fast component (24, 25, 27, 38). However, this required selecting the suitable range in advance. Importantly, the single-knee model failed to reveal the slow- and fast-component timescales.

### The slow component selectively tracked cardiac physiology

Next, we asked if the slow-component timescale that did not show functional gradient reflected potential relations with cardiac-related activity, because the spectral knees clustered over the slow component were near heart rate (mean 1.2 Hz in the present dataset; ∼1 Hz in prior work (28)) and heart rate variability can influence aperiodic spectral properties (39). We therefore tested whether cardiac physiology was selectively associated with the slow component.

At the contact level, we calculated correlations between simultaneously recorded ECG and sEEG after filtering both signals into 1-10 Hz and 10-100 Hz bands, which contained the slow and fast components, respectively (**Fig. 3a**). Contacts were classified as single knee slow, single knee fast, or double knee. In the low-frequency 1 to 10 Hz band, ECG-sEEG correlations were stronger in both classes containing a slow component than in single knee fast contacts (*p* = 0.0003 for both comparisons, bootstrap, FDR corrected; **Fig. 3b**, left). In the high-frequency 10 to 100 Hz band, correlations were lower in both classes containing a fast component than in single knee slow contacts (*p* = 0.0003 for both comparisons; **Fig. 3b**, right). Together, ECG-sEEG coupling increased with the presence of the slow component at low frequencies but decreased with the presence of the fast component at high frequencies, suggesting that the slow component was specifically associated with cardiac activity.

**Figure 3.**
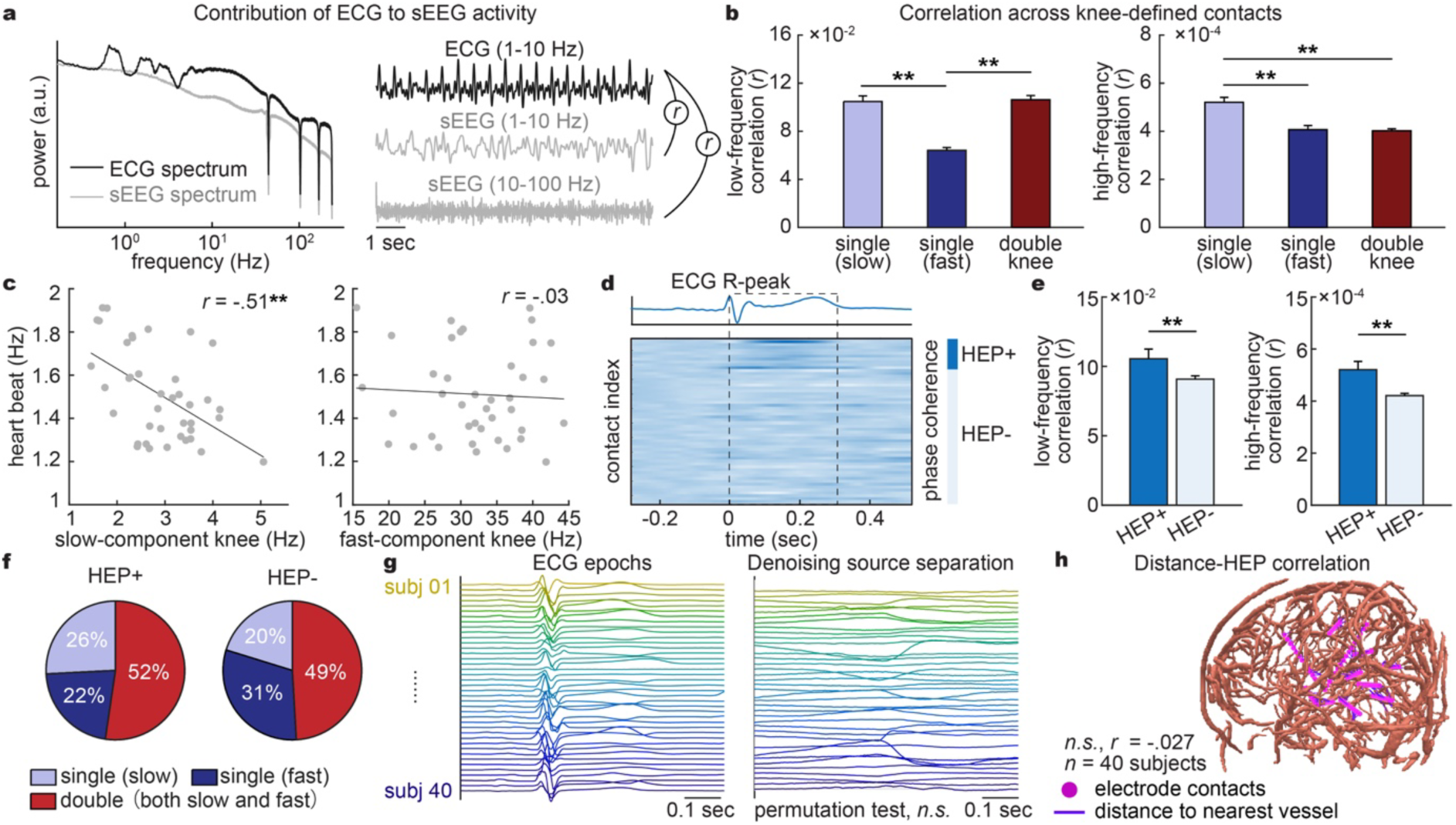
The slow component is selectively associated with cardiac physiology and heartbeat-evoked neural activity. (**a**) Analysis of ECG-sEEG coupling. Left, representative PSDs of simultaneously recorded ECG and sEEG. Right, representative ECG and band-limited sEEG signals. Coupling is quantified separately at 1-10 Hz and 10-100 Hz, the frequency ranges containing the slow and fast components, respectively. (**b**) ECG-sEEG coupling across spectral classes. Contacts are classified as single-knee contacts containing only a slow component, single-knee contacts containing only a fast component, or double-knee contacts containing both components. At 1 to 10 Hz, coupling is stronger in both classes containing a slow component than in single-knee fast contacts. At 10 to 100 Hz, coupling is stronger in single-knee slow contacts than in either class containing a fast component. Error bars denote s.e.m. across contacts. (**c**) Associations between heart rate and knee frequency across subjects. Each point represents one subject (*n* = 40); knee frequencies are summarized by the median across contacts within each subject. Heart rate is associated with the slow-component knee frequency but not with the fast-component knee frequency (two-sided Spearman’s rho). (**d**) Identification of heartbeat-evoked potentials (HEPs) in an exemplar subject. Top, ECG waveform aligned to the R peak. Bottom, intertrial phase coherence of R-peak-aligned sEEG activity from a representative subject. HEP+ contacts were identified by comparing intertrial phase coherence across sEEG epochs spanning 0 to 300 ms after each ECG R peak with a contact-specific permutation null distribution. (**e**) ECG-sEEG coupling, quantified as in (**b**), in HEP+ and HEP-contacts. (**f**) Spectral-class composition of HEP-positive and HEP-negative contacts. Pie charts show the proportions of single-knee slow, single-knee fast, and double-knee contacts within each HEP group. (**g**) Control for direct ECG leakage using denoising source separation. Left, R-peak-aligned ECG epochs from the 40 subjects. Right, the corresponding sEEG 1^st^-ranked DSS components. For each subject, no significant ECG-like components were found. (**h**) Control for local vascular pulsation. A representative reconstruction shows sEEG contacts (magenta) and subject-specific vasculature (red); distance is measured from each contact to the nearest reconstructed vessel surface. \**p* < 0.05; \*\**p* < 0.005.

At the subject level, we exploited inter-individual variation in heart rate. For each of the 40 subjects, we separately summarized slow and fast-component knees by taking the median across contacts. Slow-component knee correlated with heart rate across subjects (*ρ* = −0.509, *p* = 0.0009, two-sided Spearman’s rho), whereas fast-component knee frequency showed no such relation (*p* = 0.87). A direct component-by-heart-rate analysis further confirmed that the association differed significantly between components (*p* = 0.0023, lme model), with heart rate negatively associated with the slow component (*β* = −2.51, *p* = 0.0029) but not the fast component (*β* = −0.12, *p* = 0.88), after controlling for age and sex. The contact- and subject-level analyses therefore converged on a selective association between the slow component and cardiac physiology. These analyses established component specificity but did not determine whether the association reflected neural activity or cardiac contamination, which we examined next.

### Heartbeat-evoked neural activity linked cardiac physiology to the slow component

To distinguish these possibilities, we identified contacts with significant heartbeat-evoked potentials (HEPs) by testing whether the intertrial phase coherence of R peak aligned sEEG activity exceeded a permutation-derived null distribution (1,000 random permutations, thresholded at 95^th^ percentile, *p* < 0.05; **Fig. 3d**) (40, 41). Contacts with significant phase-locked HEPs also showed stronger ECG-sEEG amplitude coupling in the 1 to 10 Hz range than contacts without HEPs (*p* = 0.039, two-sided bootstrap, FDR corrected; **Fig. 3e**). More importantly, a slow component was present in 26% of one-knee contacts and 52% of two-knee contacts with HEPs, but was decreased to 20% and 49%, respectively, without HEPs (odds ratio = 2.014, *p* = 0.003, mixed effects logistic regression; **Fig. 3f**). Thus, the slow component was preferentially expressed at contacts showing HEPs, linking this spectral component to local responses to cardiac signals.

We next asked whether these HEPs could instead reflect direct ECG leakage or local vascular pulsation. Despite local bipolar rereferencing, we screened every denoising source separation (DSS) component from all subjects for QRS template similarity. Only two subjects showed at least one significant ECG-like component and were excluded from all analyses; no component was significant in the remaining subjects (*p* > 0.05, permutation test, FDR corrected). For illustration, **Fig. 3g** showed the first-ranked DSS component from each of the 40 subjects included in the cardiac analyses. As an independent control for local vascular pulsation, HEP magnitude was unrelated to the distance from each contact to the nearest reconstructed vessel surface (*n* = 474 contacts from 40 subjects; Pearson’s *r* = −0.027, *p* = 0.55; linear mixed effects model: *β* = −0.003, *p* = 0.10; **Fig. 3h**). Together, these controls support a heartbeat-evoked neural contribution to the slow component and argue against detectable ECG leakage or local vascular pulsation.

### A consistent dual-component temporal architecture across conscious states

We next asked whether the dual-timescale architecture identified during wakefulness was a stable property of cortical dynamics or a configuration specific to that state. We tracked spectral knees within subjects across active wakefulness, resting-state task, sleep, and anesthesia. In every state, knee frequencies retained a bimodal distribution, and a two-component model was consistently favored over a one-component model (model comparison, ΔBIC > 5.4, see Methods; **Fig. 4a**), demonstrating that the dual-component architecture persisted across marked changes in brain state. The absolute timescales nevertheless varied with state. Both the slow and fast components had their faster timescales (i.e., higher-frequency knee) during active wakefulness and slower timescales (i.e., lower-frequency knee) in less active states (all *p* < 0.012, bootstrap, FDR corrected; **Fig. 4b**).

**Figure 4.**
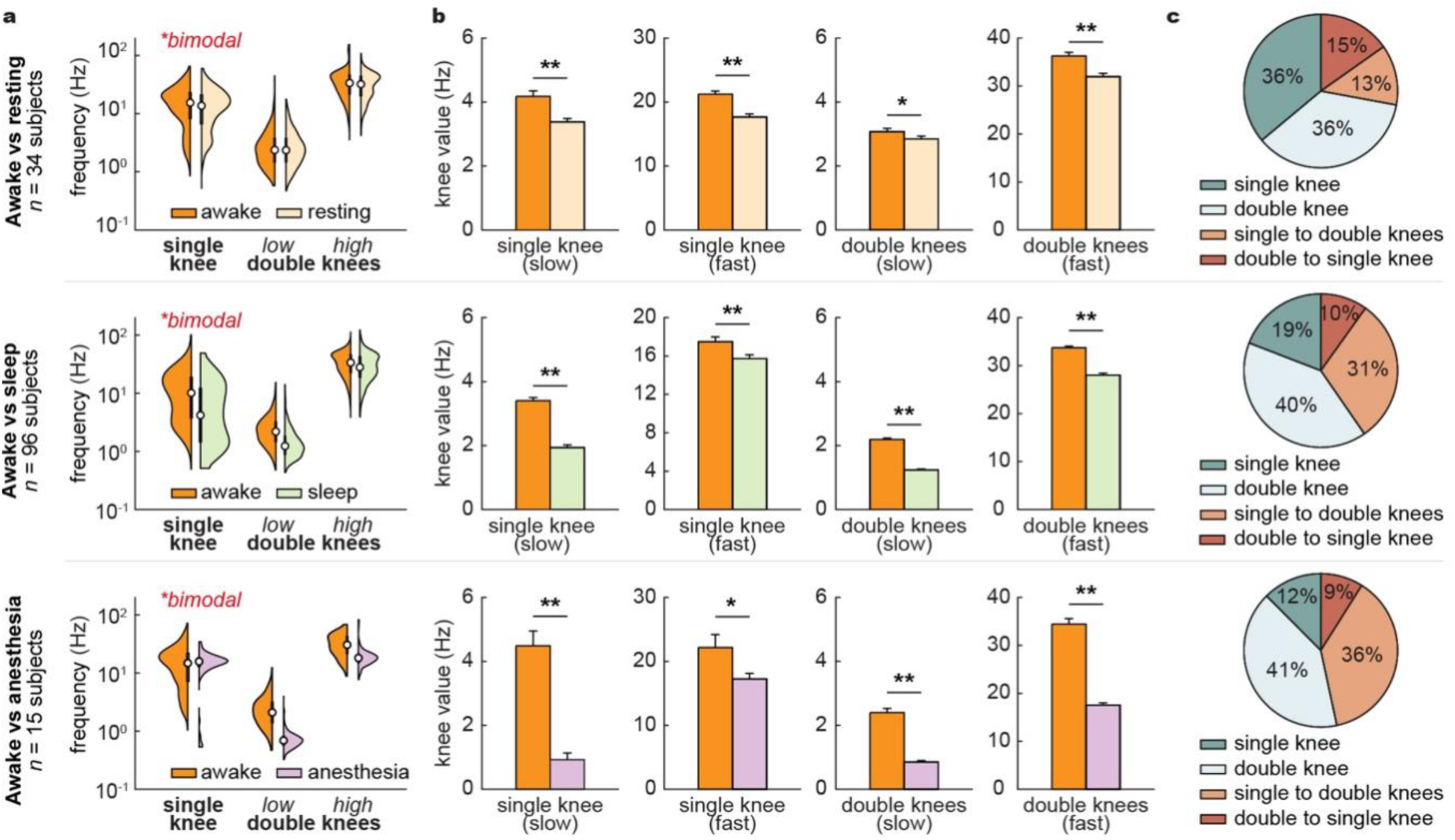
Consistent two-timescales organization across consciousness states. (**a**) Distributions of knee frequencies across consciousness states. For contacts with a single knee, a consistent bimodal organization of low- and high-frequency clusters was observed in all states. Another group of contacts with double-knee was also observed with similar low- and high-frequency clusters in all states. (**b**) Knee values change across conscious states. Both slow- and fast-component knee frequencies increase in the awake state and decrease progressively in less active states. Error bars denote s.e.m across electrode contacts. (**c**) Transitions of spectral types (i.e., contacts with one-knee or two-knee) across awake to resting, sleep, and anesthesia states. More than half of the contacts preserved their original spectral class, whereas the proportion of transitions differed by state. \**p* < 0.05, \*\**p* < 0.005.

To assess stability at individual sites, we tracked whether each contact was classified as single-knee or double-knee across states. More than half retained the same class across state transitions (**Fig. 4c**). Class changes were less frequent between wakefulness and rest, whereas transitions involving sleep or anesthesia were more frequent and showed a shift toward double-knee spectra. Similar to the coupled-process simulation in **Fig. 1b**, these transitions could reflect state-dependent changes in the interaction between persistent slow and fast processes, causing the same local system to express either one or two resolvable knees. Thus, the dual-timescale architecture remained stable across brain states, while its expression at individual contacts was state dependent.

## Discussion

Understanding the human brain’s diverse temporal infrastructures is challenging because of methodological constraints and requires high recording resolutions. By applying a novel method to large-scale human intracranial recordings, we demonstrate that the brain dynamics are built upon two dissociable temporal components. The fast component selectively encodes the gradient of timescale, whereas the slow component operates independently, coupling to the rhythms of the bodily apparatus without gradient in cortical hierarchy. This dual-component temporal architecture is intrinsic and stable across states.

Many biological systems, including neural circuits, operate across multiple interacting time scales; when these scales are sufficiently separated, the dynamics can often be decomposed into coupled fast and slow subsystems (42). The dual-timescale structure we observe is therefore broadly consistent with established biophysical frameworks (43–45) and recent studies that report stable low- and high-frequency knees in LFP spectra (32). These components form two separable, brain-wide distributions over 0.2–200 Hz rather than a single unimodal continuum. This dissociation suggests that human temporal processing is organized around two dominant regimes with distinct functional roles.

The fast component aligns with the canonical cortical hierarchy, with shorter integration in sensory regions and progressively longer integration toward association cortex, consistent with prior accounts linking the timescale gradient to sensorimotor processing and cognitive integration (11, 12, 46–48). Notably, classical evidence for this gradient was established in spiking activity, where spike-train autocorrelation statistics vary systematically along the hierarchy (9, 25), and the fast component we identify ranges at higher frequencies that are closely related to population spiking activity (49). Therefore, separating fast and slow temporal structure is necessary to faithfully link LFP dynamics to hierarchical cortical computation.

In contrast to the fast component, the slow component remains less well understood and has been variably interpreted in current models. One influential account links slow cortical dynamics to distributed long-range recurrent connectivity, in which global coupling supports slow integration beyond local fast dynamics (33). Moreover, recent work has argued that aperiodic neural activity can substantially reflect cardiac variability (39), and we find convergent evidence for this interpretation. In the current study, the slow component showed markedly stronger coupling to cardiac signals than the fast component (**Fig. 3b**). Indeed, the cardiac cycle can rhythmically modulate population-level neural activity, yielding measurable phase-dependent fluctuations in ongoing brain dynamics. These observations indicate that the slow component may capture not only distributed neural recurrence but also endogenous, physiology-linked dynamics that shape ongoing neural activity and have received less attention in neurophysiology and modeling. Together, these findings support a two-system view in which a slow, organism-level process integrates internal state and maintains homeostasis, thereby providing the context within which fast cortical computations operate (49, 50).

This discovery was enabled by overcoming a core methodological constraint. While FOOOF remains one of the most accessible tools for aperiodic parameterization, its core limitation lies in a prior assumption of a single Lorentzian function on the PSD. Importantly, for non-invasive modalities such as EEG and MEG, the single-knee assumption may be similarly misleading since a substantial proportion of second knees fall below 40 Hz (51). Whereas FOOOF provides established measures of aperiodic exponents and oscillatory peaks, the current implementation of SPLIT focuses on knee number and frequency; its segment-specific exponent estimates require further validation, and oscillations are handled separately using IRASA. Developing a unified framework that jointly quantifies multiple knees, exponents, and oscillatory activity is therefore an important direction for future work.

By transcending the oversimplified assumption of a single timescale, we establish a new analytic framework that extends from prior observations of oscillations and aperiodic exponents (26, 32, 52–55) to adopt a multi-scale lens of aperiodic and periodic neural activities. Our findings inspire a paradigm shift: the brain’s temporal organization is not dominated by one, but multiple, parallel dynamical subsystems. The parallel temporal infrastructures enable the human brain to collectively regulate the internal bodily apparatus and interact with the external world.

## Methods

### Simulated neural signals

#### Generation of the simulated PSD

To evaluate the performance of the piece-wise linear fitting procedure, we generated synthetic power spectra that contained predefined changes in spectral slope (i.e., multiple bend-knee points). These spectra were designed to avoid computational bias in estimating PSD while mimicking signals with heterogeneous timescales, such that each segment of the spectrum followed a distinct linear trend on a log-log scale. The ground-truth locations of the slope changes, as well as the slope values for each segment, were explicitly controlled and therefore known for every simulated spectrum. The simulated aperiodic PSD, *P*(*f*), was defined as:

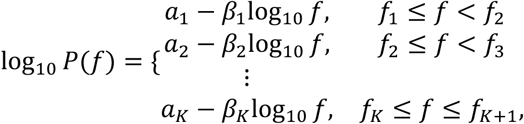

where *β_k_* denotes the ground-truth slope for segment *K* ∈ {1,2,3}, and *a_k_*ensures continuity at the boundaries. Vertices frequencies and slope values were parameterized explicitly, enabling direct comparison between ground truth and estimates by the algorithm.

White noise was added independently at each frequency and scaled by a multiplicative factor *m*, producing the final simulated spectrum:

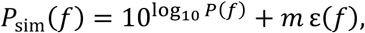

where *ε*(*f*) denotes Gaussian white noise. Noise levels spanned a predefined range to evaluate the robustness of the algorithm under varying signal-to-noise conditions.

All spectra were simulated over 0.2 to 200 Hz with 0.2-Hz resolution. For each condition defined by the number and value of knee points and noise level, a thousand spectra were generated. For each spectrum, we quantified (i) deviation between estimated and ground-truth knee values, (ii) the accuracy of recovering the correct number of knee points, and (iii) goodness of fit.

#### OU process with a single timescale

To generate a synthetic time-series of neural signals with a single intrinsic decay timescale, we simulated a one-dimensional Ornstein-Uhlenbeck process. The process satisfies the stochastic differential equation:

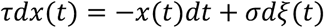

where *τ* is the decay constant, *σ* describes the strength of noise, and *ξ*(*t*) is the Brownian motion process. Numerical integration was performed using the Euler-Maruyama scheme:

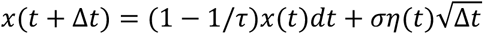

with a step size of Δ*t* = 0.001 s (i.e., 1 kHz sampling rate). Produced stationary OU signals exhibiting a single exponential decay in their autocorrelation structure.

#### OU process with double timescales

To construct synthetic signals containing two intrinsic timescales, we simulated a two-dimensional OU system and used the sum of its two components as the final observable signal. The system dynamics are given by:

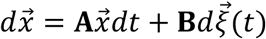

with a system matrix and a noise strength matrix:

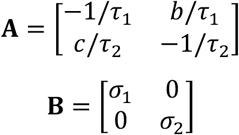

Thus, the two coordinates evolve as coupled OU subprocesses with distinct decay constants *τ*_1_ and *τ*_2_, where *b* and *c* describe the strength of coupling between two components and are set to 0 when two components evolve independently. The systems were integrated using the Euler-Maruyama update:

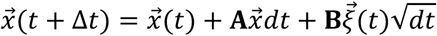

Each simulation began from [0,0]*^T^* and ran for 15 minutes at 1 kHz. The final signal was defined as the linear sum:

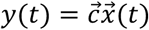

Where 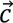 is the summation vector. This yields a process whose autocorrelation is the superposition of two exponential decays with timescales *τ*_1_ and *τ*_2_.

#### OU process with oscillation(s)

To generate signals exhibiting both stochastic decay and oscillatory features, we extended the OU model by introducing a sinusoidal component to the OU signals described above. After simulating the OU system and forming the mixture signal *x*_OU_(*t*), we added one or more oscillatory components of the form:

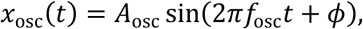

and final signal was constructed as:

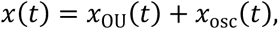

Taken together, all three kinds of OU synthetic time series combining single or double timescales, with or without oscillations, were used to evaluate the algorithm’s ability to distinguish oscillatory components from aperiodic structure, and to test robustness under mixed periodic-stochastic conditions.

### Empirical recordings and neural signals

#### Subjects

This study analyzed fully de-identified intracranial EEG data originally collected for clinical monitoring in patients with medically intractable epilepsy at the Guangdong Sanjiu Brain Hospital (Guangzhou, China) and Shenzhen Second People’s Hospital (Shenzhen, China). The present research involved secondary analysis of legacy clinical data and did not require any additional data collection or patient contact. All procedures were approved by the Research Ethics Committee of the two hospitals and the Institutional Review Board of New York University Shanghai.

A total of 185 subjects (70 females; ages 18-52 years, mean age 29 years) were included. We analyzed three subsets of subjects, each providing recordings from four physiological states to enable within-subject comparisons: (i) awake dataset (136 subjects), consisting of subjects with recordings in awake state; (ii) awake-sleep dataset (96 subjects), consisting of subjects with recordings during awake and asleep; (iii) awake-resting dataset (34 subjects), consisting recordings during subjects were awake and in task-free resting state; and (iv) awake-anesthesia dataset (15 subjects), consisting of subjects with recordings during awake and general anesthesia.

#### Physiological Recording States

Awake state: Awake recordings were collected when patients were fully conscious and responsive during routine clinical monitoring. Patients were not given behavioral instructions and were free to engage in spontaneous everyday activities (e.g., conversation, phone use). A 15-minute segment was extracted. In addition, for analyses relating the low-frequency knee to heart rate, we selected extended awake segments (30 mins to 4 hours) that contained concurrent ECG recordings.

Resting state: Resting-state recordings were obtained while patients were awake and instructed to relax without performing any task. Patients were asked to keep their eyes open or fixate on a visual point, remain still, and refrain from speaking or moving. No sensory stimulation or cognitive tasks were presented. A 5-minute resting-state segment was used for analysis.

Sleep state: Sleep recordings were acquired during natural overnight clinical monitoring. For each patient, a 15-minute segment was randomly selected from natural sleep.

Anesthesia state: Recordings when subjects were under general anesthesia were obtained during clinically indicated procedures in which anesthetic agents (e.g., propofol or sevoflurane, depending on clinical requirements) were administered by the clinical team. No task or stimulation was presented during anesthesia maintenance. A 15-minute segment was extracted for analysis.

#### sEEG data collection and preprocessing

Stereo-EEG was recorded with clinical depth electrodes (Sinovation, China) implanted in the brain parenchyma. Signals were amplified with a 256-channel Neuvo system and sampled at 2,000 Hz. Each electrode comprised 8-16 cylindrical contacts (0.8 mm diameter; 2 mm length; 1.5 mm inter-contact spacing). Electrode trajectories were planned with the manufacturer’s ROSA software by co-registering preoperative MRI and CT. Postoperative CT acquired with electrodes in place was then co-registered to the preoperative MRI to verify implantation accuracy relative to the planned paths. The number of electrodes and placement were determined solely based on clinical purposes by the neurologists to localize epileptogenic regions as part of the presurgical evaluation.

All analyses were performed in custom MATLAB scripts (56, 57). Continuous sEEG was visually screened by a neurologist (J.C.) and analysts (Y.L. and H.M.) to identify epileptiform and artifactual activity. Data were resampled to 1,000 Hz; a narrow notch (48-52, 98-102, 148-152, and 198-202 Hz) removed line noise and its harmonics. No additional broadband high-pass filtering was applied before spectral estimation. Signals were re-referenced with a local bipolar montage by differencing adjacent contacts on each shaft (the terminal contact was omitted). Contacts that were labeled as white matter or as N/A by anatomical parcellation were excluded, and outlier contacts were removed based on abnormally high mean amplitude.

#### Structural image and co-registration

For each patient, contact localization and anatomical labeling were performed in Brainstorm (58). The T1 image was segmented with CAT12 and normalized to MNI152 space using SPM (59). The postoperative CT was co-registered to the MRI with SPM12, and contacts were manually identified on the co-registered image, and their coordinates were extracted in native MRI space. To obtain standardized anatomical locations, each contact’s native-space coordinates were transformed into MNI152 space using the deformation fields estimated during MRI normalization. This procedure allowed an MNI coordinate for each contact for cortical gradient analysis (see Cortical annotation).

#### ECG data processing and heartbeat-related statistical analyses

Concurrent ECG recordings were first notch filtered to remove line noise. R-peaks were detected using an automated peak-detection procedure and manually quality-checked to remove spurious detections and ectopic beats. Mean cardiac frequency was calculated for each participant by taking the reciprocal of each R-R interval and averaging these values across accepted heartbeats, expressed in Hz (**Fig. 3c**).

To quantify ECG-sEEG coupling, sEEG were further filtered into a low-frequency band (1-10 Hz) and a higher-frequency band (10-100 Hz), and Pearson correlations were computed between the band-limited ECG (1-10 Hz) and each contact’s band-limited sEEG over the continuous recordings (**Fig. 3b**).

Intertrial phase coherence was used to identify contacts showing sEEG activity consistently phase-locked to heartbeats. Phase estimates were obtained using Fourier transform and epoched relative to each ECG R peak. Statistical testing was restricted a priori to the 0 to 300 ms post-R interval. For each contact, the observed intertrial phase coherence within the prespecified time window was compared with a null distribution generated from 1,000 permutations, in which phase values were randomly shuffled. Contacts whose observed intertrial phase coherence exceeded the permutation threshold were classified as HEP positive; all remaining contacts were classified as HEP negative (**Fig. 3d**).

To test for direct ECG leakage, DSS was applied separately within each participant. The subject-specific ECG QRS template and the R-peak-aligned waveform of each DSS component were standardized, and their similarity was quantified as Spearman correlation within the prespecified QRS interval. For each DSS component, significance was assessed against a subject specific null distribution generated by circularly shifting the DSS time series relative to the ECG R peaks by a random nonzero lag 1,000 times. Empirical *p* values were corrected across DSS components within each subject using FDR. Subjects containing at least one component with significant QRS template similarity after correction were excluded from all analyses. Two subjects who had positive significant DSS components were excluded.

#### MR angiography

All vascular analyses began with raw magnetic resonance angiography (MRA) images. Vessel enhancement and segmentation were performed in 3D Slicer using a vesselness-based filter to improve contrast of tubular vascular structures against background tissue. Subject-specific vascular masks were generated and were co-registered in SPM to each participant’s T1-weighted anatomical image using a rigid-body transformation, ensuring spatial alignment across vascular, anatomical, and electrode coordinate spaces. Finally, three-dimensional surface models of the vasculature were reconstructed from the aligned masks via the marching cubes algorithm. Vertex coordinates of each model were converted into world coordinates using the appropriate affine transform, enabling consistent distance measurements between electrode locations and nearby vascular structures.

### Timescale estimate pipeline

#### Aperiodic spectrum estimation using IRASA

Oscillatory components were removed using the Irregular-Resampling Auto-Spectral Analysis procedure, which separated the broadband aperiodic spectrum from narrowband rhythmic peaks through non-integer resampling (60). For each sEEG electrode contact, continuous data were divided into 70-s sliding windows with 50% overlap (**Table 1**), and the signal was resampled using a set of multiplicative factors h and 1/h (e.g., ℎ = 1.1, 1.2, …, 1.9), stretching and compressing oscillatory components while leaving the fractal 1/f background unchanged. Averaging the spectra across all resampling factors canceled out the oscillatory components, yielding a fractal (aperiodic) spectrum. IRASA was applied over 0.01-400 Hz (due to algorithmic requirement (55)) and the fractal spectrum was taken forward for knee fitting.

**Table 1.**
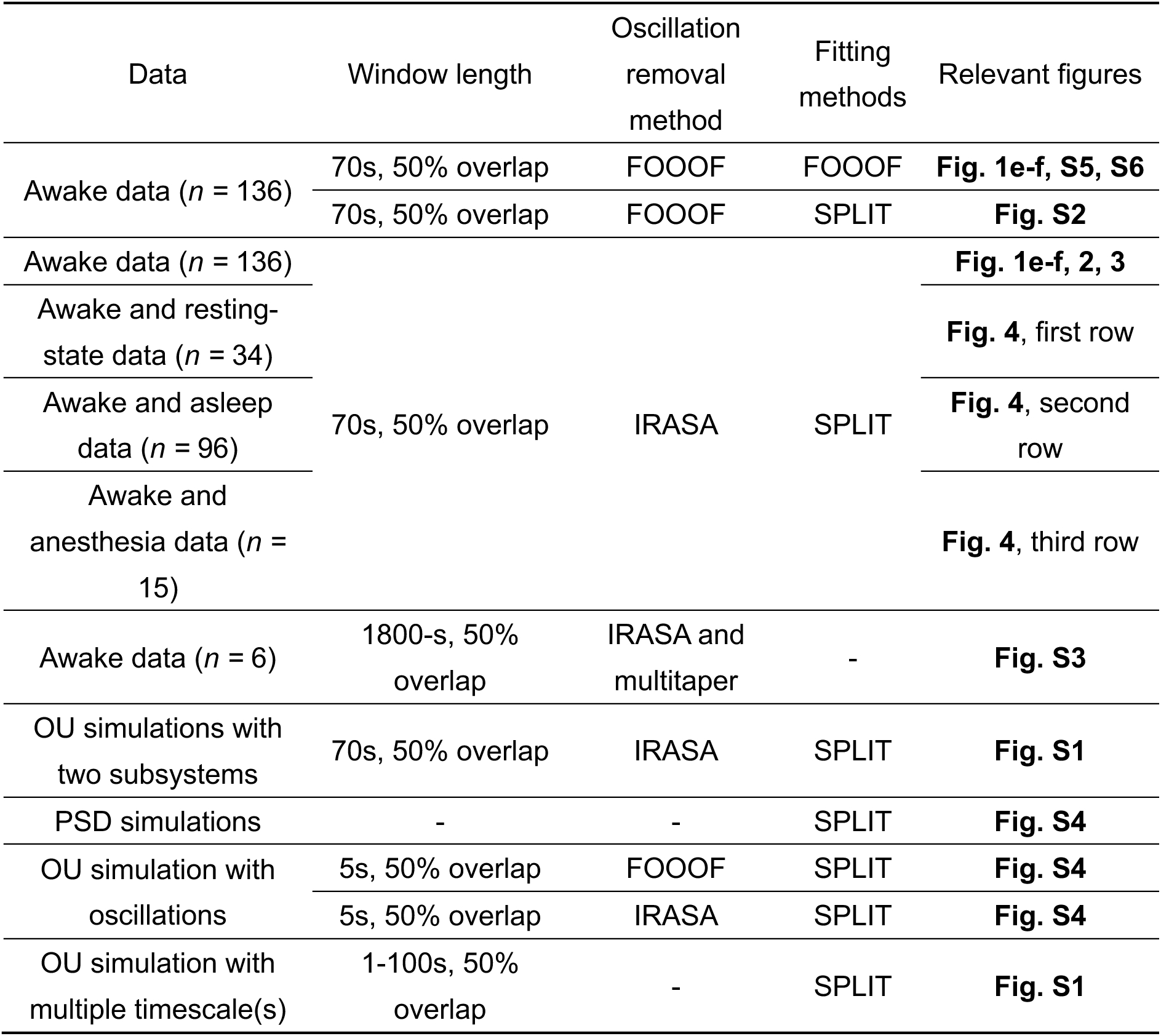
Summary of datasets and analysis configurations.

#### SPLIT (spectral piecewise-linear inference of timescale) fitting

To identify multiple spectral timescales in the aperiodic spectrum, we modeled PSD in log-log space, where 1/f-like spectra became approximately piecewise linear. In this domain, each linear segment corresponds to a distinct power-law regime, and the transitions between segments defined characteristic timescales. Because the frequency sampling was highly non-uniform in log space, with dense sampling at high frequencies and sparse sampling at low frequencies, direct fitting would overweight the high-frequency range. To ensure equal contribution across frequencies, the log-transformed PSD was linearly interpolated onto an evenly spaced grid of 1,000 log-frequency points spanning the full fitting range (0.2-200 Hz). This procedure preserved the 1/f curvature while standardizing sampling density for segmentation. We carefully evaluated grid densities ranging from 100 to 2,000 points, with steps of 100. Knee estimates improved with increasing grid density and had effectively converged by 1,000 points we used, differing subtly from estimates obtained at higher densities (**Fig. S7a**). Note that increasing grid density would substantially increase computational cost with negligible gain in estimation accuracy.

Subsequently, for each contact, we fitted a piecewise linear model with K segments (*K* = 1, …, 4), corresponding to zero to three spectral knees:

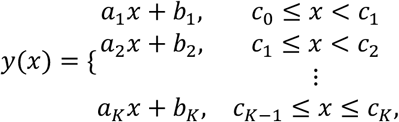

This parameterization enforced continuity between adjacent segments. For a given K, breakpoint locations and regression coefficients were jointly optimized by differential evolution as implemented in pwlf function. The local aperiodic exponent of each segment was defined as the negative of its slope, and each breakpoint was converted to a knee frequency as *f*_knee_ = 10*^c^*^j^ Hz. Model complexity was controlled using a penalty defined as:

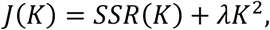

Here, SSR(K) was the residual sum of squares over the 1,000-point log-frequency grid, and λ controlled the trade-off between goodness of fit and model complexity. Smaller values imposed weaker penalties on additional segments, whereas larger values favored simpler models. We calibrated λ exclusively using simulations with known ground-truth knee numbers (**Fig. S7b**). Candidate values from 0.1 to 1.0 were evaluated in increments of 0.1. The simulations included zero-knee spectra, single-knee spectra with knee frequencies from 1 to 10 Hz in 0.5-Hz increments, and double-knee spectra with low-frequency knees from 2 to 8 Hz in 0.5-Hz increments and high-frequency knees from 20 to 32 Hz in 1-Hz increments. For each ground-truth configuration, 100 independent realizations were generated and fitted. Performance was evaluated in two stages. First, knee-number accuracy was defined as the proportion of realizations for which SPLIT recovered the correct number of knees. Second, among realizations with the correct knee number, knee-frequency error was calculated as 100 × | knee_fit_ − knee_true_| / knee_true_ and averaged across realizations; errors for double-knee spectra were averaged across the low- and high-frequency knees. Knee-number accuracy was the primary criterion because frequency error was defined only when the correct model was recovered. At λ = 0.5, recovery was simultaneously near ceiling for zero-, one-, and two-knee spectra, while conditional knee-frequency errors remained near their lower plateau. We therefore fixed λ at 0.5 for all empirical spectra.

Eventually, for each spectrum, we searched the discrete candidate set K and selected the model that minimized the penalized objective J(K). The selected model was refitted with pwlf to obtain the final breakpoints and segment slopes. Note that we only retained spectra in which successive segments became progressively steeper with increasing frequency; spectra violating this ordering were treated as unclassified. This automated procedure was applied to all spectra. Y.L. and H.M. carefully inspected the fitted spectra as a quality control step; this inspection did not alter model selection or parameter estimation. For transparency and reproducibility, all contact spectra and fitted curves obtained across the tested regulatization weights are available in the data repository (see *Data availibility*).

#### Comparator methods

Welch and multitaper methods: For comparisons using conventional spectral estimates, Welch spectra were computed from the same preprocessed signals using 70 s windows with 50% overlap and averaged across windows for each contact. To examine spectral structure over longer durations, recordings from six subjects were additionally analyzed using multitaper spectra computed from 1,800 s windows with 50% overlap and compared with IRASA estimates (**Fig. S3** and **Table 1**). These alternative spectral estimates were used only for control analyses.

FOOOF: For comparison with models with single-knee assumption, we fit each power spectrum using the FOOOF algorithm, which parameterizes the aperiodic component using a Lorentzian 1/f function and Gaussian peaks. FOOOF was applied across simulations and empirical data analysis using the following parameters: knee mode, a maximum of three oscillatory peaks (max_n_peaks = 3), peak width limits of 1-12 Hz (peak_width_limits=[1, 12]), a minimum peak height of 0.2 (min_peak_height = 0.2), and a peak-detection threshold of 2 (peak_threshold = 2). All spectra were fit over the 0.2-200 Hz frequency range. FOOOF-based oscillation removal was also used only for comparison, using the above-mentioned parameters. For cross-method validation, oscillatory peak components identified by FOOOF were removed from the original spectrum, and SPLIT was applied to the resulting peak-removed aperiodic spectrum.

#### Assignment of knees to slow and fast components

To assess whether the data exhibited a unimodal or bimodal distribution, we adopted a model-based approach using Gaussian mixture models (**Fig. 2a-b**, and **4a**). The data were first log-transformed, after which Gaussian mixture models with one to four components were fitted using the expectation-maximization algorithm. To ensure robust parameter estimation, each model was fitted with multiple random initializations, and covariance regularization was applied to avoid numerical instability. Model selection was performed using the BIC, which balances goodness of fit against model complexity. If the model with two components yielded the lowest BIC, indicating that a bimodal distribution provided the most parsimonious explanation of the data.

### Cortical hierarchy annotation

To place local timescale estimates within established macroscale organization, we examined their distribution across three widely used cortical atlases: the seven intrinsic functional networks (35), the principal unimodal-transmodal functional gradient (10), and the HCP T1w/T2w ratio map, a myelin-sensitive proxy of anatomical cortical hierarchy (37). For the network analysis, contacts were grouped according to their assigned functional network. For the gradient analysis, contacts were assigned to ten equally populated bins according to the gradient value of their cortical location. Knee frequencies were averaged separately for the slow and fast components within each network or gradient bin. For the myelin analysis, we used the group-average T1w/T2w ratio map from the WU-Minn Human Connectome Project S1200 release provided in HCP-MMP1.0 surface space. Following Gao et al. (25), the median T1w/T2w value was calculated across surface vertices within each parcel and subsequently z-scored across the parcels included in the analysis. Knee frequencies were averaged across contacts within each corresponding parcel, separately for the slow and fast components and for single-knee and double-knee spectra, and then related to the parcelwise T1w/T2w values.

### Statistical analysis

Contact-level analyses used bootstrap or linear mixed-effects models. Parcelwise analyses treated cortical parcels as the sampling units. Benjamini-Hochberg false discovery rate (FDR) correction was applied to control for multiple comparisons.

#### Bootstrap test

Bias corrected and accelerated bootstrap resampling (61) was used to compare spectral classes, HEP groups, and brain states was used to compare the difference between conditions (**Fig. 3b**, **3e** and **Fig. 4b**). In the bootstrap procedure, participants were resampled 10,000 times with replacement. For each comparison, we calculated the proportion *A* of bootstrap samples in which the mean knee value in one condition was greater (or smaller) than in the other condition. All tests were two-sided, with formula to obtain the p-value: *p* = (2 ⋅ min(*A*, 1 − *A*) ⋅ 10,000 + 1)/10,001.

#### Correlation analysis

Pearson correlation was used to quantify associations between knee frequencies estimated by SPLIT and FOOOF (**Fig. 1e-f**) and between knee frequency and cortical hierarchy (**Fig. 2d-f**). Because at least one of the two variables was not normally distributed, the association between mean heart rate and median knee frequency across subjects was assessed using a two-sided Spearman rank correlation (**Fig. 3c**).

#### Linear mixed effect model

To complement the correlation analyses and account for repeated measurements across channels and subjects, we further performed linear mixed-effects modeling. For cortical hierarchy, component, gradient, and their interaction were included as fixed effects, with subject and channels nested within subject as random intercepts: *knee* ∼ *component* ∗ *gradient* + (1|*subject*) + (1|*subject*: *c*ℎ*annel*) (**Fig. 2d-f**). Models were fitted separately for each hierarchy measure and for single- and double-knee spectra. For the cardiac analysis, we used *knee* ∼ *component* ∗ ℎ*eart rate* + *age* + *sex* + (1|*subject*) to directly test the component-by-heart-rate interaction (**Fig. 3c**). For the relation between HEP amplitude and Euclidean distance to vessel, we used the equation: HEP ∼ distance + (1 | subject) + (1 | subject: channel) (**Fig. 3h**). All models were fitted using MATLAB’s fitlme function.

#### Across-state spectral transitions

Only contacts recorded in both states of a given comparison were included (**Fig. 4c**). Single-knee and double-knee retention and transition proportions were calculated within each subject. Because the awake-to-rest, awake-to-sleep, and awake-to-anesthesia analyses included different subject cohorts, transition rates were not compared inferentially across cohorts.

## Data availability

The de-identified sEEG datasets supporting the findings of this study are available at http://github.com/[…]. Readers interested in accessing the raw dataset may contact the lead author. Data access will be granted to named individuals in accordance with ethical guidelines for the reuse of clinical data, subject to the completion of a formal data-sharing agreement and institutional approval.

## Code availability

The analysis codes supporting the findings of the study are available at our GitHub repository: https://github.com/[…]. The SPLIT algorithm code is openly available in both MATLAB and Python versions with supporting documents at https://github.com/[…].

## Acknowledgement

We sincerely thank Youcun Zheng, Ruoyu Wu, Mingxuan Fang, Xiaofeng Zeng, Yuling Huang, and Peiyu He for assisting in data preparation. This study is supported by the National Natural Science Foundation of China 32271101 (X.T.), the Brain Science and Brain-like Intelligence Technology - National Science and Technology Major Project No. 2022ZD0204600 and the Natural Science Foundation of China Grant No. 31771147 (Z.L. and L.T.), and the Program of AI-Driven Initiative to Promote Research Paradigm Reform and Empower Disciplinary Advancement by Shanghai Municipal Education Commission (SMEC), Program of Introducing Talents of Discipline to Universities, Base B16018, and NYU Shanghai Boost Fund to X.T.

## Author contributions

All authors initiated and designed the study. Y-H.L. and H.M. conceived of and coded the algorithm. Y-H.L., H.M., Z.L., and Q.L. contributed code and tested the algorithm against real and simulated data. J.C. and Y.C. collected data. Y-H.L., H.M., Q.L., and Y-D.L. analyzed data. L.T, Z.X. and X.T. provided overall supervision and guidance throughout the project. All authors contributed to the manuscript.

## Declaration of interests

The authors declare no competing financial interests.

## Notes

### Competing Interest Statement

The authors have declared no competing interest.

### Summary of Updates

We have uploaded revised manuscripts based on the Reviewers's comments.

