## Supplementary figures for "Neural Architectures of Slow and Fast Dynamics in the Human Brain"

**Supplementary Document**

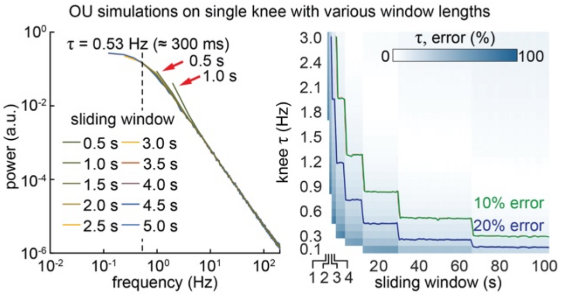


**Fig. S1. Window duration constrains the recovery of low-frequency spectral knees.** Left: Welch power spectra from a 15-min Ornstein-Uhlenbeck process with a single ground-truth knee at 0.53 Hz, corresponding to τ = 1/(2πf_knee) ≈ 300 ms. Spectra were estimated using Hamming windows ranging from 0.5 to 5.0 s with 50% overlap. Colors indicate window duration, and the vertical dashed line marks the ground-truth knee. Short windows distorted the spectral curvature around the knee and displaced the apparent bend toward higher frequencies. Right: Recovery error across ground-truth knee frequencies from 0.1 to 3.0 Hz and sliding window length from 1 to 100 s. For each combination of knee frequency and window length, 100 OU realizations were independently generated and fitted. Each cell shows the mean absolute relative error in recovered knee frequency, calculated as 100 × | $knee_{fit}$ – $knee_{true}$ | / $knee_{true}$. Green and blue contours mark mean errors of 10% and 20%, respectively. A 70-s window constrained the mean error to below 20% for knee frequencies at or above approximately 0.2 Hz, supporting its use in the empirical analyses.

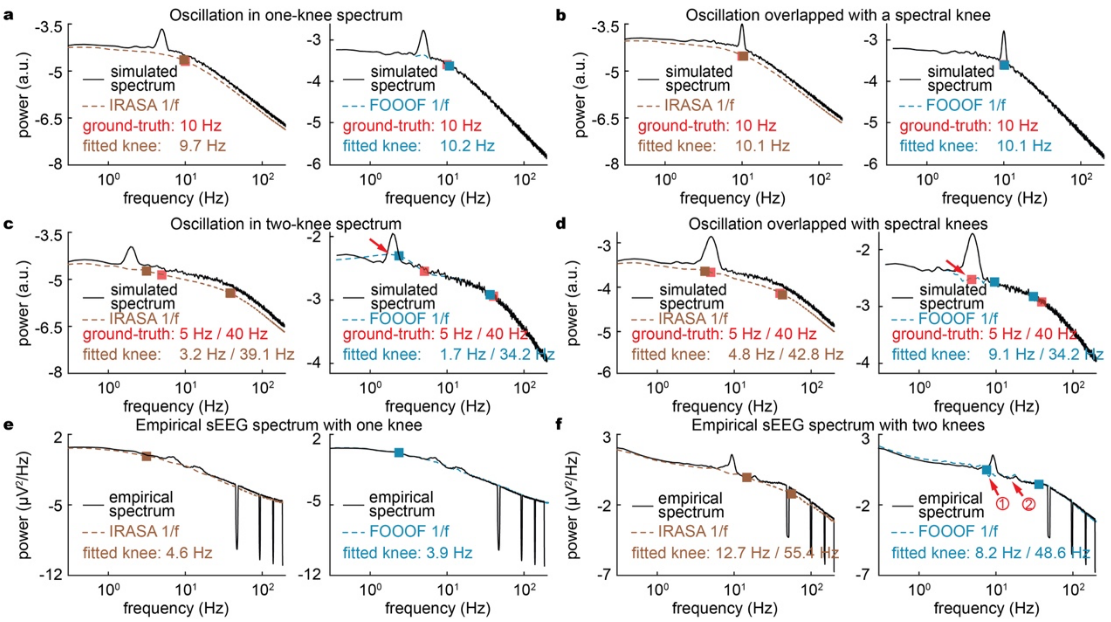


**Fig. S2. Oscillation removal based on a single-knee model can distort spectra containing multiple knees.** (**a-b**) Simulated spectra containing one ground-truth knee at 10 Hz and a narrowband oscillation positioned away from the knee (**a**) or centered on it (**b**). Black curves show the simulated spectra. Brown dashed curves show the fractal spectra estimated by IRASA, and blue dashed curves show the spectra obtained after removal of oscillatory peaks identified by FOOOF. Red squares indicate the ground-truth knees, whereas brown and blue squares indicate knees subsequently recovered by SPLIT and FOOOF, respectively. Both approaches recovered the single knee closely. (**c-d**) Simulated spectra containing ground-truth two knees at 5 and 40 Hz, with an oscillation positioned near the low-frequency knee (**c**) or overlapping it (**d**). IRASA more closely preserved the underlying spectral curvature and recovered both knees near their ground-truth locations. FOOOF-based peak removal introduced local deviations around the oscillation and displaced the subsequent knee estimates, as indicated by the arrows. (**e-f**) Representative empirical sEEG spectra containing one knee (**e**) or two knees (**f**), identified by two investigators Y.L. and H.M.. The two approaches produced similar estimates for the single-knee spectrum. In the double-knee example, FOOOF-based peak removal altered the local spectral curvature and shifted both knee estimates; numbered arrows mark the corresponding local deviations. Ground-truth and recovered knee frequencies are reported within each panel. All spectra are shown on logarithmic frequency and power axes.

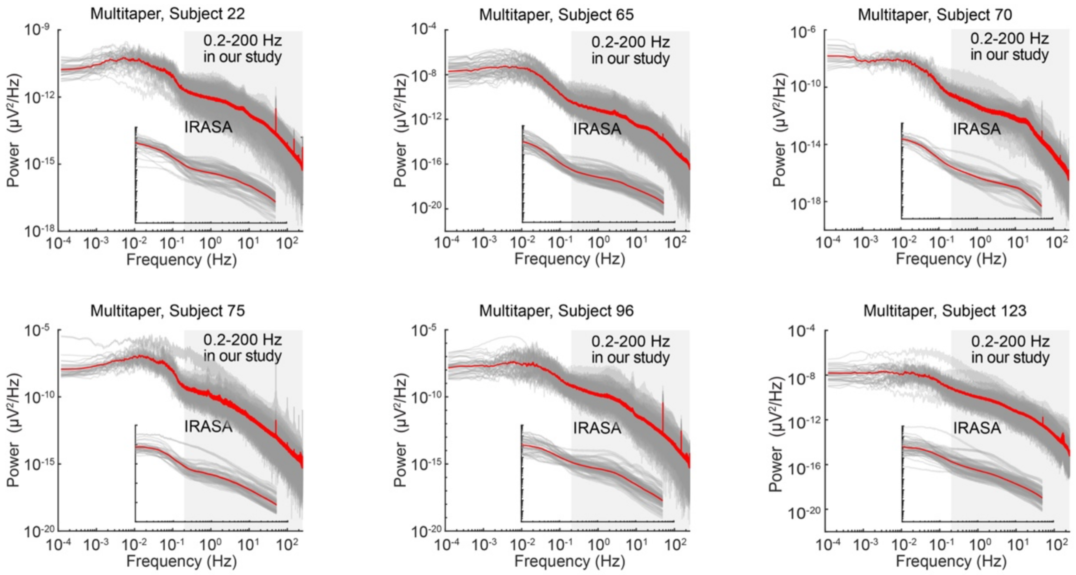


**Fig. S3. Spectral curvature below 0.2 Hz lies outside the prespecified fitting range.** Multitaper power spectra were computed from extended recordings in six subjects using 1,800-s windows with 50% overlap. Gray curves represent individual contacts, and the red curve shows the across-contact mean for each subject. Shading marks the prespecified 0.2 to 200 Hz range submitted to SPLIT. Insets show the corresponding IRASA-derived fractal spectra using the same color convention (gray shading also marks the 0.2 to 200 Hz range in these plots). Pronounced and subject-dependent curvature was present below 0.2 Hz, but this frequency range was not included in knee estimation in the present study.

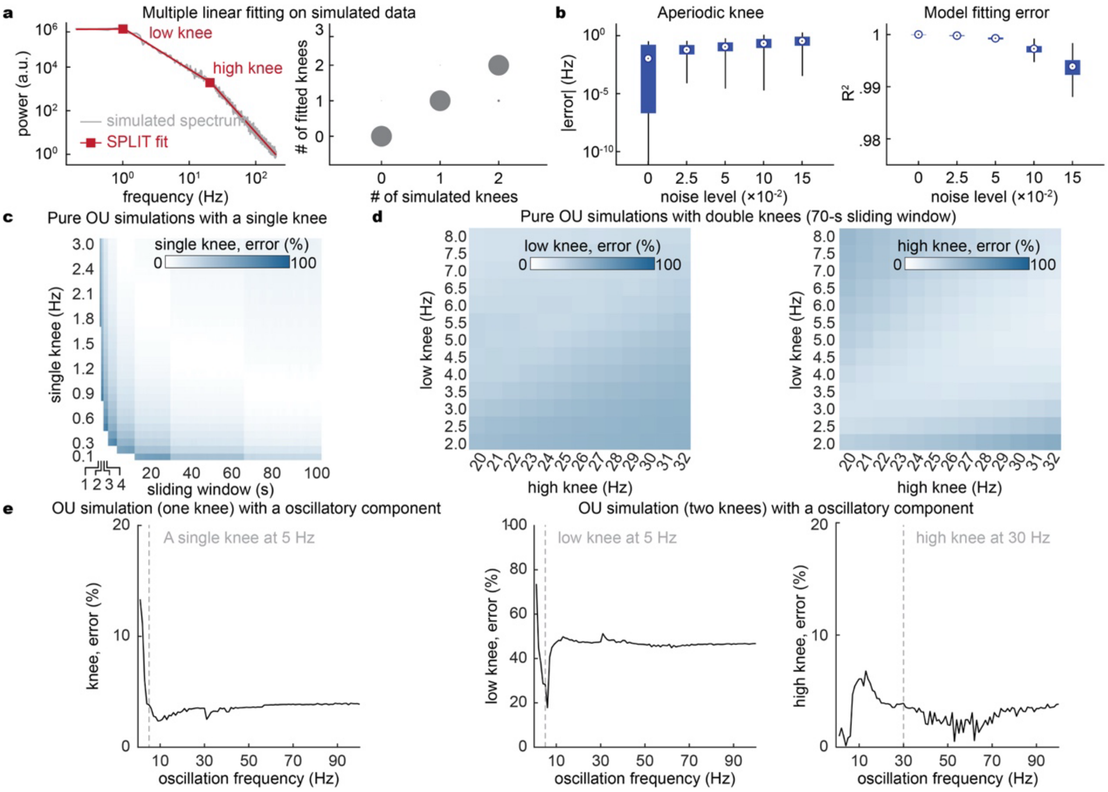


**Fig. S4. Ground-truth simulations characterize the performance and limits of SPLIT.** (**a**) Recovery of knee number from idealized piecewise-linear spectra. Left, representative spectrum containing two ground-truth knees (gray) and the corresponding SPLIT fit (red); squares indicate the recovered knees. Right, recovered knee number plotted against ground-truth knee number for spectra containing zero, one, or two knees. Circle area indicates the relative frequency of each outcome. Most outcomes lie on the diagonal, indicating recovery of the correct knee number. (**b**) Robustness to additive spectral noise. Absolute knee-frequency error (left, logarithmic scale) and goodness of fit, quantified by $R^2$ (right), are shown across increasing noise amplitudes. Knee-frequency errors remain small, while $R^2$ remains near 1 throughout the tested range. Box-and-whisker plots summarize the distributions across simulations. (**c**) Recovery of a single knee from OU signals. Mean absolute relative error in knee frequency is shown as a function of the ground-truth knee frequency, from 0.1 to 3.0 Hz, and window duration, from 1 to 100 s. Each condition is averaged across 100 independently generated and fitted realizations. Lower-frequency knees require longer windows for accurate recovery. (**d**) Recovery of two knees from OU signals using 70-s windows. Mean absolute relative error is shown separately for the low-frequency knee (left) and high-frequency knee (right), across combinations of low knees from 2 to 8 Hz and high knees from 20 to 32 Hz. Each condition is averaged across 100 independently generated and fitted realizations. The high-frequency knee is recovered more precisely, whereas estimation of the low-frequency knee is more sensitive to the underlying knee configuration. (**e**) Effect of a superimposed oscillation. Left, relative error for a single-knee OU signal with a ground-truth knee at 5 Hz as oscillation frequency is varied. Middle and right, relative errors for the low-frequency knee at 5 Hz and high-frequency knee at 30 Hz in a two-knee OU signal. Vertical dashed lines indicate the corresponding ground-truth knee frequencies. Single-knee and high-frequency-knee estimates vary modestly with oscillation frequency, whereas the low-frequency knee in the two-knee signal remains less precise. Relative error in (**c**) to (**e**) is calculated as 100 × | $knee_{fit}$ – $knee_{true}$ | / $knee_{true}$.

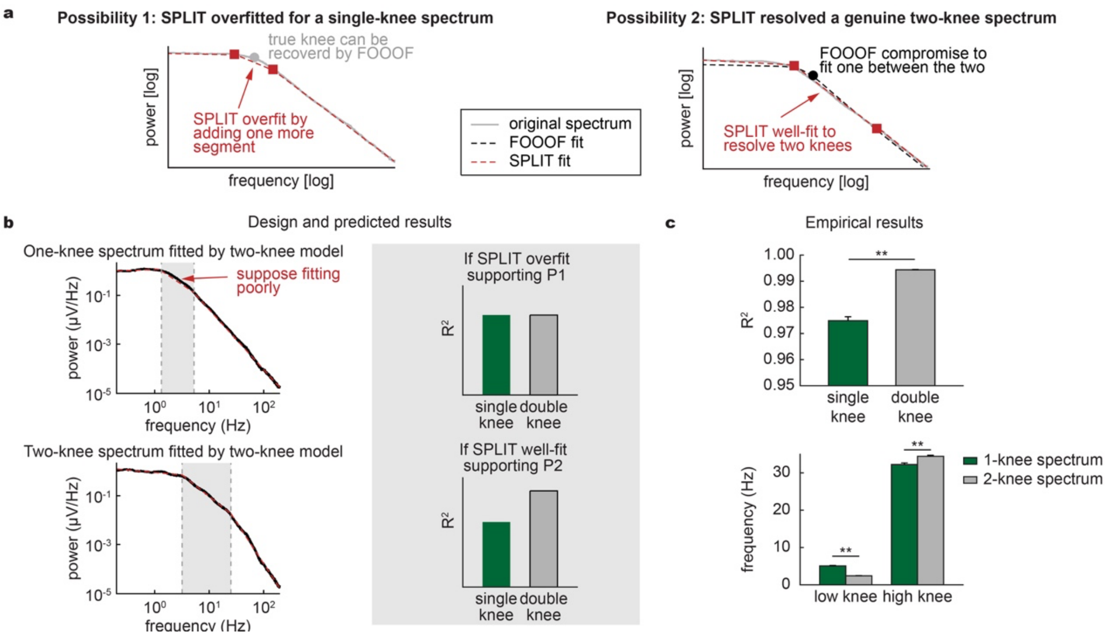


**Fig. S5. A linear intermediate segment distinguishes double-knee spectra from an overfit single bend.** (**a**) Competing explanations. Left: One possibility is that SPLIT may have divided one broad bend into two knees, with FOOOF recovering the underlying single knee. Right: The other possibility is that SPLIT may have resolved two distinct bends, with the single-knee FOOOF model returning a compromise between them. (**b**) Discriminating test. Left: a two-knee model was forced onto single-knee spectra (top) and compared with the same model fitted to double-knee spectra (bottom). The intermediate segment (shading) crosses the curved transition in a single-knee spectrum but follows an approximately linear regime between two distinct knees. Right, the overfitting hypothesis predicts similar intermediate-segment $R^2$ values in the two groups, whereas the two-bend hypothesis predicts a higher $R^2$ in double-knee spectra. (**c**) Observed results. Top, intermediate-segment $R^2$ was higher in double-knee spectra. Bottom, when two knees were forced onto single-knee spectra was significantly differed from those knees in two-knee spectra. These results argued against subdivision of a single bend and supported two distinct knees. Bars show means, and error bars indicate s.e.m. $p = 0.0002$ for all comparisons, two-sided bootstrap tests with FDR correction. $^{**}p < 0.005$.

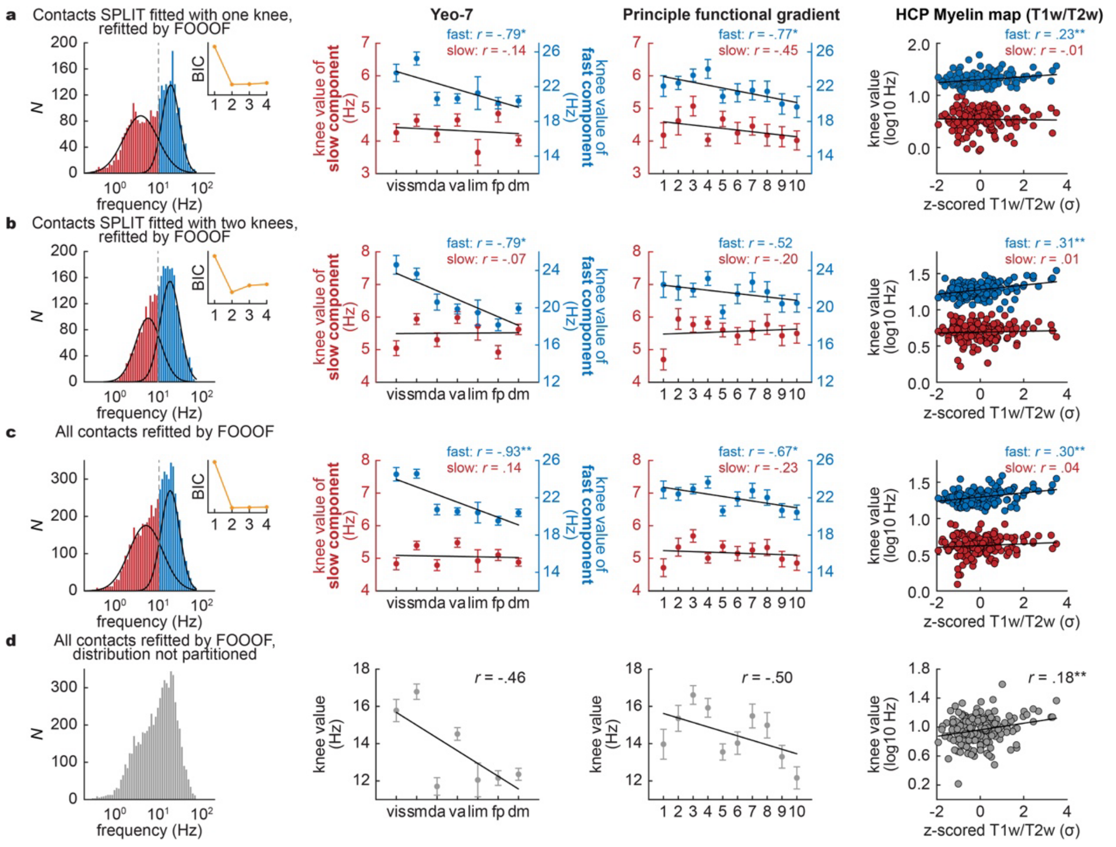


**Fig. S6. FOOOF estimates reveals the cortical hierarchy of the fast component.** (**a**) FOOOF estimates from contacts classified as single-knee by SPLIT. (**b**) FOOOF estimates from contacts classified as double-knee by SPLIT. (**c**) FOOOF estimates from all contacts combined. In (**a-c**), Gaussian mixture models containing one to four components were fitted to log-transformed knee frequencies. Histograms show estimates assigned to the slow (red) and fast (blue) clusters. Black curves show the fitted component densities, and the vertical dashed line marks their intersection. Insets show BIC values, with lower values indicating better model support. The remaining columns show knee frequencies across the Yeo seven-network organization, across ten equally populated bins of the principal functional connectivity gradient, and as a function of the parcel-wise HCP T1w/T2w myelin map. The fast cluster was significantly associated with the Yeo network hierarchy and cortical myelination in all three contact sets, and with the principal functional gradient in the single-knee and pooled analyses. No corresponding association was detected for the slow cluster. (**d**) The same FOOOF estimates from all contacts analyzed without population-level separation. Pooling the estimates removed the significant associations with both functional hierarchy measures and attenuated, but did not eliminate, the association with cortical myelination. Error bars indicate s.e.m. across contacts within each network or gradient bin. Points in the myelin analyses represent cortical parcels. Pearson's r values are reported in each panel; significance was assessed using one-sided tests with FDR correction. Network abbreviations are vis, visual; sm, somatomotor; da, dorsal attention; va, ventral attention; lim, limbic; fp, frontoparietal; and dm, default mode. $^*p < 0.05$; $^{**}p < 0.005$.

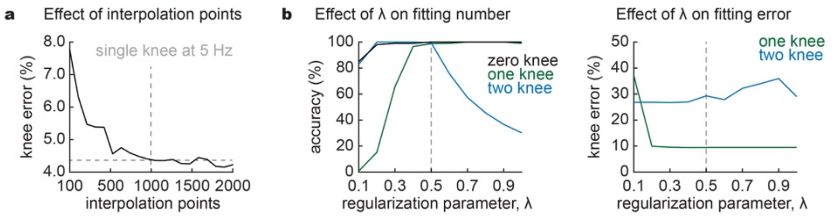


**Fig. S7. The selection of interpolation density and model-complexity regularization in SPLIT.** (**a**) Effect of interpolation grid density. A simulated spectrum containing a single ground-truth knee at 5 Hz was interpolated using grids ranging from 100 to 2,000 equally spaced log-frequency points. Mean absolute relative error in the recovered knee decreased as grid density increased and changed minimally beyond approximately 1,000 points. The vertical dashed line marks the selected grid density of 1,000 points; the horizontal dashed line marks the corresponding error. (**b**) Selection of the regularization weight λ using simulations with known knee numbers. Left, percentage of realizations in which SPLIT recovered the correct number of knees at each λ. Lower λ values reduced the recovery of zero- and one-knee spectra, whereas higher values reduced the recovery of two-knee spectra. Right, mean absolute relative knee-frequency error, calculated only among realizations in which the correct knee number was recovered. For double-knee spectra, errors were averaged across the low- and high-frequency knees. Frequency error was not defined for zero-knee spectra. The vertical dashed line marks λ = 0.5, which provided near-ceiling knee-number accuracy across all three classes while maintaining conditional frequency errors near their lower plateau. This value was fixed for all empirical analyses.
